# Single cell trajectory analysis of human pluripotent stem cells differentiating towards lung and hepatocyte progenitors

**DOI:** 10.1101/2021.02.23.432413

**Authors:** Chaido Ori, Meshal Ansari, Ilias Angelidis, Fabian J. Theis, Herbert B. Schiller, Micha Drukker

## Abstract

Understanding the development of human respiratory tissues is crucial for modeling and treating lung disorders. The molecular details for the specification of lung progenitors from human pluripotent stem cells (hPSCs) are unclear. Here, we use single cell RNA-sequencing with high temporal resolution along an optimized differentiation protocol to determine the transcriptional hierarchy of lung specification from human hPSCs and map out the underlying single cell trajectories. We show that Sonic hedgehog, TGF-*β* and Notch activation are required in an *ISL1*/*NKX2-1* trajectory that gives rise to lung progenitors during the foregut endoderm stage. Induction of *HHEX* marks an alternative trajectory at the early definitive endoderm stage, which precedes the lung trajectory and generates a major hepatoblast population. Moreover, neither KDR+ nor mesendoderm progenitors are apparent intermediate states of lung and hepatic lineages. Our hierarchical multistep model predicts mechanisms leading to lung organogenesis, and creates a basis for studying early human lung development, as well as hPSC based disease and drug research.

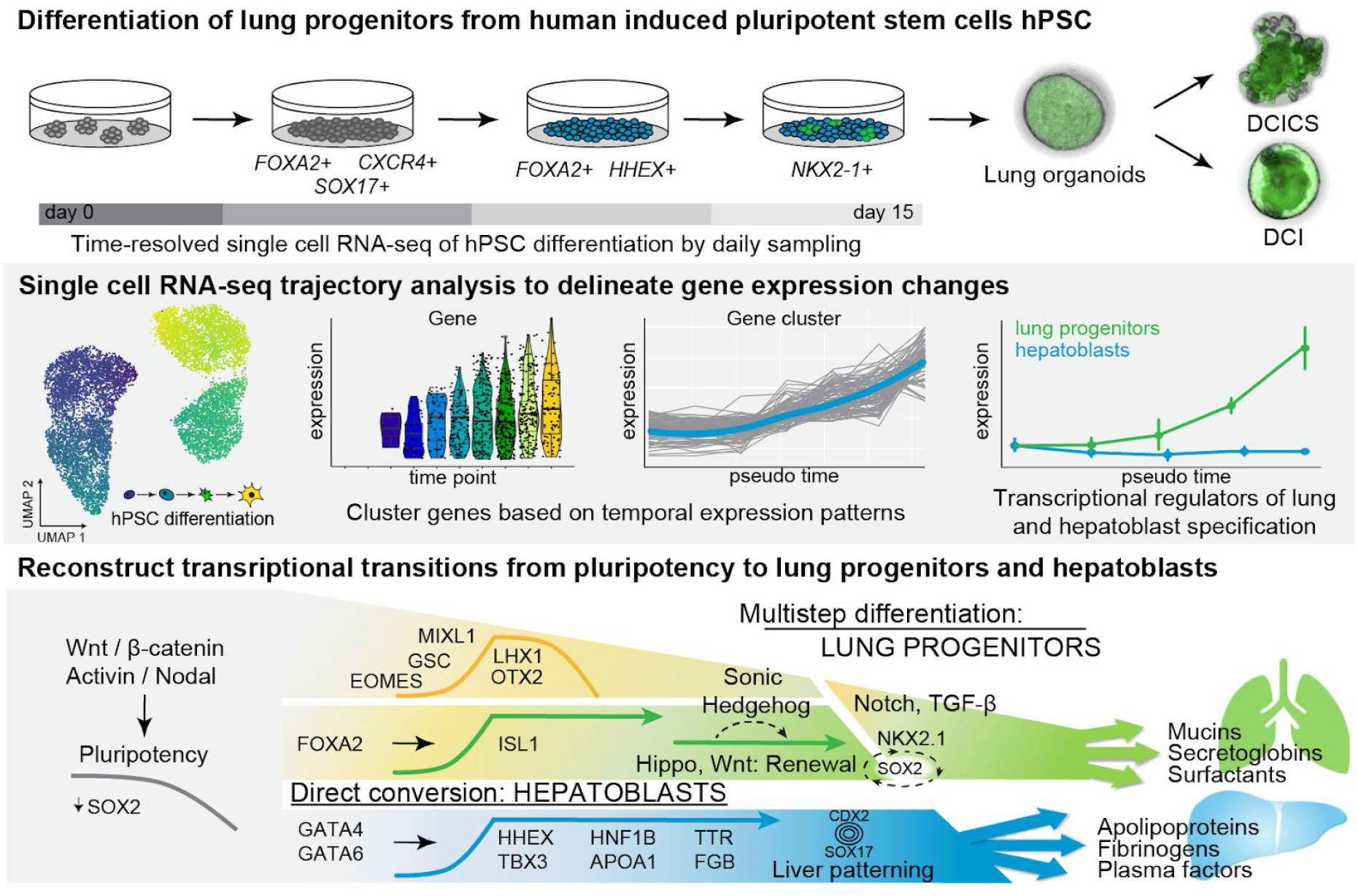

## INTRODUCTION

A primary goal of stem cell biology is to understand how the intricate regulation of cellular differentiation by cascades of gene regulatory networks and signaling pathways gives rise to the structure and function of organs. This knowledge is fundamental to human pluripotent stem cell (hPSC)-based approaches that model human development and congenital pathologies *in vitro*, and to devise regenerative therapies. The elucidation of molecular mechanisms governing foregut endoderm (FE) formation is of particular interest for biomedical research, as the FE gives rise to several organs, including the lung and the liver, that are relevant for disease modelling, drug screening assays and regenerative applications. Various definitive endoderm (DE) progenitors, and the genes and pathways that regulate their specification, still need to be better characterized in order to effectively control the production of cells that represent endodermal tissues *in vitro*, such as epithelial cells of the airways and hepatocytes.

Single cell genomics is the method of choice for analyzing cell-cell heterogeneity and cellular differentiation trajectories in development (Argelaguet et al., 2019; Griffiths et al., 2018; Nowotschin et al., 2019; Pijuan-Sala et al., 2019). Single cell RNA sequencing (scRNA-seq) applied to differentiation protocols of hPSCs can be thus used to understand aspects of human development from *in vitro* studies. For instance, the molecular “roadmap” governing the development of functional human islet beta cells has been characterized recently using this approach by applying scRNA-seq time-series experiments (Veres et al., 2019). While embryological studies in mice have been instrumental in revealing the major stages and genes involved in foregut development, insight into the early steps of human lung specification is at large pending. Recent studies have taken important steps in this direction by characterizing signaling pathways that regulate the differentiation of hPSC-derived NKX2-1+ progenitors into alveolar epithelial type 2 (AT2) cells (Hurley et al., 2020). NKX2-1 is a transcription factor (TF) that is essential for lung and thyroid development in the foregut (Kimura et al., 1996), and hence expression of this gene serves as a hallmark to characterize hPSC-derived progenitors that can give rise to functional lung cells (Chen et al., 2017; Hawkins et al., 2017; Longmire et al., 2012). Nevertheless, the intermediates states and mechanisms prompting the commitment of NKX2-1+ lung progenitors during endoderm differentiation have not been characterized in detail. In this regard, it is not well understood which mechanisms lie upstream of NKX2-1, and why mixtures of cells that express NKX2-1 and hepatic markers such as FGB are being produced by current state of the art differentiation protocols (Hawkins et al., 2017).

Current protocols of lung progenitor differentiation from hPSC utilize a stepwise application of signaling cues that are known to be important in lung development. Treatment of hPSCs by Activin / Nodal in conjunction to activation of Wnt/*β*-catenin signaling promotes the differentiation of DE, followed by transient inhibition of endogenous Activin / Nodal and BMP signaling by “dual SMAD inhibition”, which promotes further commitment of the FE (Green et al., 2011). In the last step Wnt/*β*-catenin is reactivated in conjunction to RA, BMP4 and FGF10 treatment, resulting in the formation of NKX2-1+ progenitors (Green et al., 2011; Huang et al., 2014). NXK2-1+ progenitors have been shown to mimic early lung organogenesis as they can produce alveolar and bronchiolar spheroids that express proteins involved in respiratory functions including surfactants (Gotoh et al., 2014; Jacob et al., 2017; Konishi et al., 2016; McCauley et al., 2017).

Differentiation protocols of hepatic lineages and lung progenitors are similar in the initial activation of Activin / Nodal pathway to specify the DE. However, contrary to lung differentiation protocols, hepatic induction involves treatment by FGF2 or FGF2+BMP4 in the DE phase (Hannan et al., 2013). In the second stage, some protocols utilize SMAD inhibition similarly to lung differentiation protocols (Green et al., 2011), while others utilize HGF (Carpentier et al., 2014). The disparities in the early induction of DE raise questions about the origin and identity of the hepatic cells that are neighboring to NKX2-1+ progenitors in lung differentiation protocols. Moreover, in this regard it is not resolved whether hepatic KDR+ progenitors (Goldman et al., 2013) or mesendoderm precursors (Burtscher and Lickert, 2009) are transient states in lung progenitor differentiation protocols.

The stalk and epithelial tips of human fetal lungs have recently been characterized by RNA sequencing (Nikolić et al., 2017), and a cell atlas for the adult human lung has been created by scRNA-seq (Travaglini et al., 2020; Vieira Braga et al., 2019). How NKX2-1+ progenitors are being formed during development remains an open question. Using time-series single cell RNA-seq we model the hierarchy of gene expression changes along a trajectory from hPSCs to NKX2-1+ lung progenitors. Our model illustrates how timed inputs of Activin A/Nodal, Wnt/*β*-catenin, hedgehog, and the TGF-*β* pathways promote lung progenitor differentiation. We discover a novel early role for Notch signaling in this process. Surprisingly, we also found that the emergence of hepatoblasts can take place directly from early DE, prior to the formation of FE, when Activin / Nodal and Wnt/*β*-catenin signaling are at their peak (Whitsett, 1998). We characterize molecular details of this branching event that specifies lung progenitors versus hepatoblast identity. The transcriptional roadmap established here thereby provides an important resource for understanding the origin and abnormal development of the human respiratory system and hepatic lineages.

## RESULTS

### Human PSCs differentiate in parallel to lung progenitors and phenotypically early liver cells

In the mouse embryo, the formation of the primitive streak (PS) marks the stage where pluripotent cells commit themselves to the precursors of the germ layers including the DE. The differentiation of DE in this process relies on the induction of TFs including *Sox17, Foxa2* and *Eomes*, primitive streak TFs including *Mixl1, Goosecoid* (*Gsc*), and the down-regulation of pluripotency genes (Brown et al., 2011; Teo et al., 2011; Vallier et al., 2009; Vincent et al., 2003). The TFs that subsequently promote the anteriorization of the primitive gut tube leading to the induction of *NKX2-1*, are less characterized both in the mouse model and the differentiation of hPSCs. To investigate mechanisms underlying the formation of lung foregut precursors in human development, we optimized a differentiation protocol that utilizes step wise activation and inhibition of Activin / Nodal and Wnt/*β*-catenin signaling (Hawkins et al., 2017; Konishi et al., 2016), with supplementation of sonic hedgehog (SHH) and FGF10 at the second stage, as these cues are essential in foregut and lung bud development (Bellusci et al., 1997; Litingtung et al., 1998; Motoyama et al., 1998) (**Fig. 1a**).

**Figure 1:**
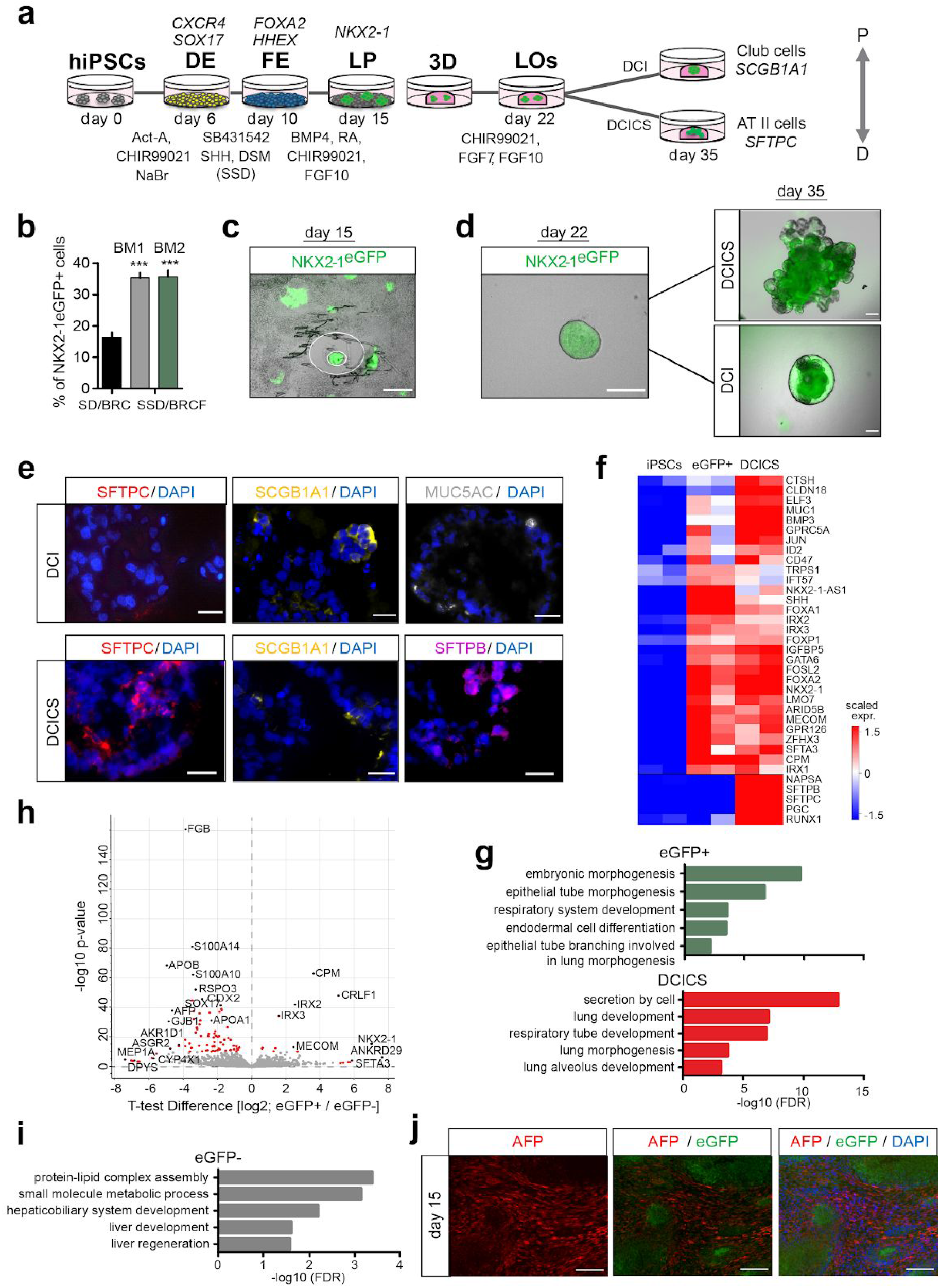
Differentiation of lung progenitors from human induced pluripotent stem cells. **(a)** A schematic illustration of the differentiation protocol used in this study to produce eGFP+ lung progenitor cells, and spheroids and organoids (on right). **(b)** Quantification of the amount of eGFP+ cells by flow cytometry on day 15 of differentiation; conditions with and w/o combined FGF10+SHH treatment in defined basal media DMEM/F12 and IMDM (FE-BM1 and BM2 respectively). Bars represent mean ± SD, n = 3 biological replicates, ***p ≤ 0.0001 unpaired two-tailed t-test. **(c**,**d)** Fluorescence microscopy of day 15 NKX2-1eGFP+ cell sectors (**c**), and a representative day 22 spheroid produced by embedding a colony in Matrigel, and the further growth of spheroids (Scale bar: 200 µm) observed on day 35 with treatments as indicated **(d). (e)** Immunofluorescence staining of day 35 DCI and DCICS spheroid-organoids for SFTPC, SFTPB, SCGB1A1 and MUC5AC (scale bar: 20μm). **(f)** A heatmap displaying the expression of developmental markers and genes important for the function of the lung based on the bulk RNA-Seq analysis of the indicated conditions (p-value < 0.05, scale displays normalized log2 expression). **(g)** Gene Ontology (GO) term enrichment analysis of differentially expressed genes in eGFP+ and DCICS spheroid-organoids relative to undifferentiated *NKX2-1*^*eGFP/+*^ cells (p-value < 0.05, GO term FDR < 0.05). **(h**,**i)** A volcano plot showing differentially expressed genes comparing the eGFP+ and eGFP-sorted populations on day 15 **(h)**, and the corresponding GO terms of the eGFP-population **(i). (j)** Immunostaining for AFP (liver marker), imaging of eGFP+ cells (lung progenitors), and DAPI staining on day 15 of differentiation (scale bars: 100 µm). DE -definitive endoderm; FE -foregut endoderm; LP -lung progenitors; LOs -lung organoids.

To monitor the appearance of lung progenitors we used a human induced PSC (hiPSC) line integrated with the *eGFP* gene downstream to the endogenous promoter of *NKX2-1* (*NKX2-1*^*eGFP/+*^) (Olmer et al., 2019). Our protocol assessment showed that *SOX17*, FOXA2, and cell surface markers that are associated with DE, namely, CXCR4, CKIT, EPCAM were markedly up-regulated at day 6 of the differentiation protocol (**Suppl. Fig. 1a,b**). The formation of eGFP+ progenitors became apparent between days 13-15, and their amount significantly increased by supplementation of SHH and FGF10, which also improved the overall cell survival (**Suppl. Fig. 1c**). We compared two types of basal media, namely, DMEM/F12 and IMDM (BM1 and BM2 as described in experimental procedures), and found superior activity of IMDM with 100% success rate and an average efficiency of approximately 35% NKX2-1+ cells located in sectors of eGFP expression with this protocol (**Fig. 1b,c and Suppl. Fig. 1c, d**). Since *NKX2-1* is expressed in the fetal lung, as well as in thyroid and forebrain progenitors (Kimura et al., 1996), we analyzed the expression of *PAX8* and *PAX6* which are representative markers of the latter tissues. The low levels of these markers on day 15 indicated that the *NKX2-1* signal represented the formation of lung progenitors but not the other tissues (**Suppl. Fig. 1e**).

Next, we analyzed the developmental potential of putative eGFP+ lung progenitors using spheroid 3D culture assays. We embedded clusters of eGFP+ cells from day 15 of differentiation in Matrigel and supplemented the media with CHIR99021 (CHIR), FGF7, and FGF10 (**Fig. 1a**), in order to promote the proliferation of lung progenitors in suspension culture (Hawkins et al., 2017). This led to the outgrowth of the spherical structures, which tripled in size within 7 days and maintained expression of eGFP (**Fig. 1d and Suppl. Fig. 1f,g**). Subsequent treatment by dexamethasone, cAMP and IBMX (DCI), which promote maturation of the fetal lung (Ballard et al., 1997; Gonzales et al., 2002), induced expression of genes and proteins that are associated with club cells and goblet cells in the proximal region of the lung, namely, SCGB1A1 and MUC5AC (**Fig. 1e and Suppl. Fig. 1f-h**). When Wnt/*β*-catenin pathway was activated by CHIR, the spheroids developed branches (DCIC), and when combined with inhibition of TGF-*β* by SB431542 (DCICS), the spheroids grew substantially larger to an average size of 1.6 mm by day 35. They also exhibited branches and markers of alveolar epithelial type II cells, namely, SFTPC and SFTPB, and lower expression of *SCGB1A1* and *MUC5AC* compared to DCI (**Fig. 1e and Suppl. Fig. 1f-h**).

To characterize the developmental stage, the differentiation potential of NKX2-1+ progenitors, and the basis of the heterogeneity in eGFP expression, we sequenced the mRNAs of undifferentiated cells, sorted eGFP+ and eGFP-populations and DCICS organoids (**Suppl. Fig.1i,j**). Genes that have been implicated in the formation of respiratory epithelial cells in the lung, namely, *FOXA2, FOXA1, FOXP1*, and *NKX2-1* (Shu et al., 2007; Wan et al., 2005), were highly expressed in the eGFP+ cells and DCICS organoids (**Fig. 1f**). GO term analysis furthermore revealed enrichment of genes involved in embryonic respiratory lung morphogenesis and alveolar development in eGFP+ progenitors and DCICS organoids (**Fig. 1g**). The organoids additionally exhibited markers that are characteristic to early and late branching morphogenesis and differentiation of the distal lung, including *RUNX1, MUC1, SFTPC, SFTPB, CLDN18* and *NAPSA* (**Fig. 1f**), and their expression pattern closely resembled published transcriptomes of the alveolar tips of human fetal lungs, but less the stalk of fetal lungs (**Suppl. Fig. 1k,l**).

Analysis of eGFP-cells revealed that the eGFP-NKX2-1-population expressed fetal liver genes, including apolipoproteins (e.g. *APOA1* and *APOB*) fibrinogen (*FGB*), plasma protein alpha fetoprotein (*AFP*) (Carpentier et al., 2014; Hannan et al., 2013), and GO term analysis revealed enrichment of fetal liver development, regeneration and metabolism categories (**Fig. 1 h,i**). To confirm the coexistence of lung progenitors and hepatocytes on day 15 of the differentiation protocol, we analyzed the cultures by immunostaining of AFP and eGFP. This showed mutually exclusive expression of eGFP and AFP, and presence of lung progenitors and hepatoblasts in separate clusters (**Fig. 1j**). Taken together, we concluded that co-induction of lung and liver fates took place during the differentiation of lung progenitors from hPSCs, raising the question of the mechanisms that drive the mutually exclusive specification of these lineages from FE.

### Time-resolved single cell RNA-seq of hPSC differentiation

To chart a comprehensive map of the transcriptional states underlying the differentiation of lung progenitors and hepatoblasts in parallel, we performed a 16-day time-series scRNA-seq using the Drop-Seq workflow (Macosko et al., 2015). Single cell suspensions were processed daily, resulting in the analysis of a total of 10,667 cells that were used for downstream analysis (**Fig. 2a and Suppl. Fig. 2a, Supplementary Table 2**). First, we used Uniform Manifold Approximation and Projection (UMAP) and Partition-based Graph Abstraction (PAGA) analysis (Wolf et al., 2019) to project the gene expression data in two dimensions and performed an assessment of the connectivity of the obtained cell clusters (**Fig. 2b,c and Suppl. Fig. 2a-c**). This revealed three major domains in the high dimensional gene expression manifold corresponding to the three stages of the differentiation protocol (**Fig. 2b, c and Suppl. Fig. 2a**).

**Figure 2:**
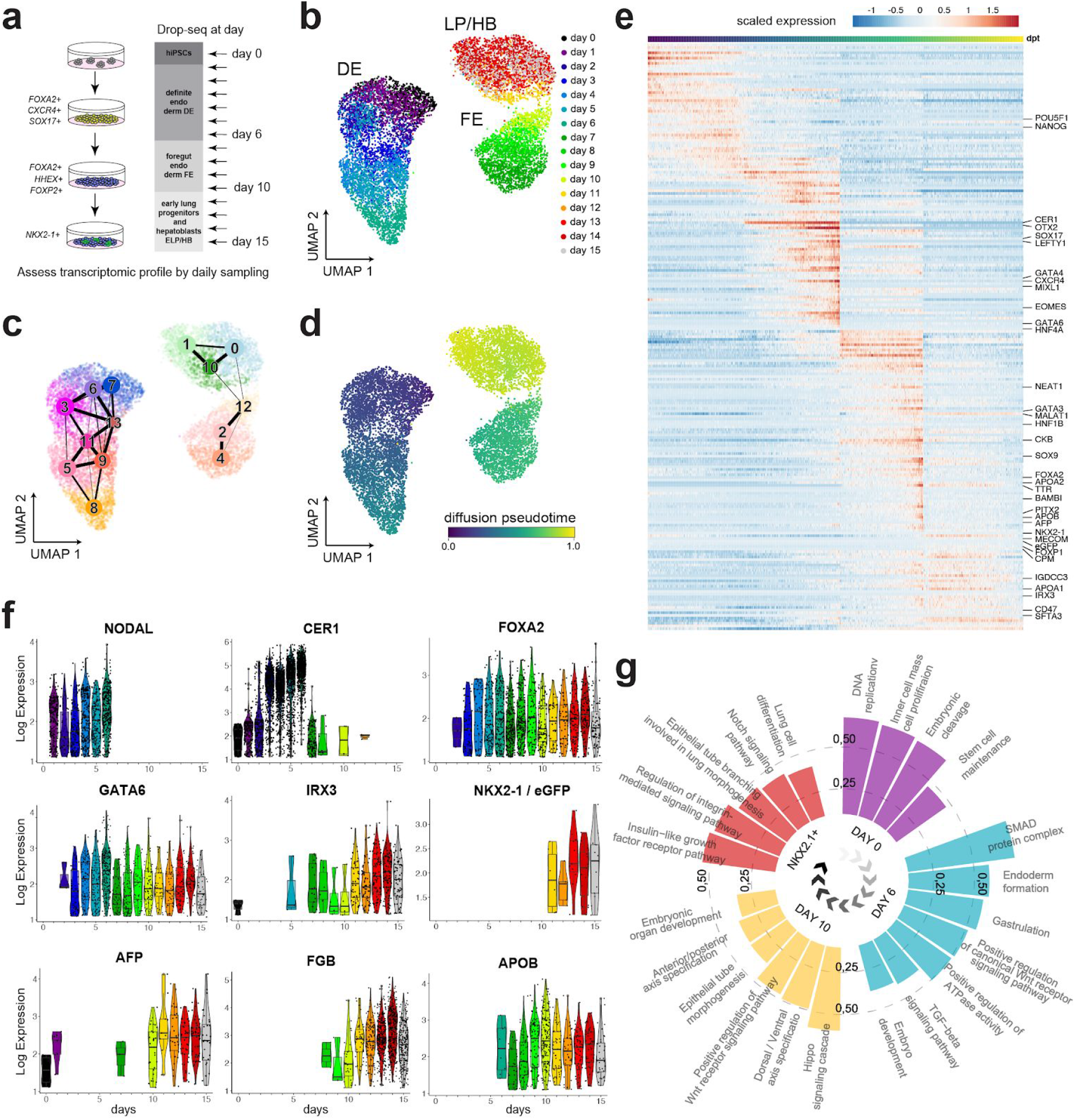
Time-resolved analysis of hPSC differentiation using scRNA-seq shows distinct hierarchy of gene expression changes. **(a)** A schematic illustration of daily sampling for scRNA-seq applied to the first 15 days of the lung progenitor differentiation protocol. **(b)** Time of sampling is color coded on the UMAP projection of scRNA-seq transcriptomes. **(c)** The connectivity of distinct louvain clusters as determined by graph abstraction (PAGA) is shown on the UMAP projection. **(d)** UMAP colored by the stage-wise inferred pseudotime; darker regions (towards blue) indicate starting points and brighter colored regions (towards yellow) ending points. **(e)** A heatmap illustrating peaks of gene expression ordered by the pseudotime of differentiation. Selected developmental markers of the lung and the liver are displayed. **(f)** Violin plots of stage specific marker genes are shown for the real time points of sampling in days. **(g)** Circular bar chart showing enriched gene categories taken from Uniprot Keywords, GO terms and KEGG pathways corresponding to the pluripotent cells, DE, foregut and NKX2-1+ progenitors (FDR <5%).

To model the dynamics of gene expression along the entire trajectory we used pseudotime inference (**Fig. 2d**), which was in a good agreement with the timepoints of sampling (**Suppl. Fig. 2d**). Unsupervised hierarchical clustering of the genes along the pseudotime ordering revealed stage specific markers, including TFs expressed in undifferentiated PSCs e.g. *POU5F1* (*OCT4*) and *NANOG*; DE factors including e.g. *SOX17, MIXL1, EOMES, CER1, CXCR4*, and *LEFTY1*; TFs characteristic for the foregut e.g. *FOXA2*, and *FOXP1* (Shu et al., 2007); and genes important for the formation of lung progenitors e.g. *NKX2-1* and *IRX3* coming up in the latest stage *(Kimura et al., 1996;van Tuyl et al ., 2006)* (**Fig. 2e and Suppl. Fig. 2b, e, f, Supplementary Table 4**). The pseudotime was consistent with consecutive expression of DE, foregut and lung progenitor markers, e.g. *NODAL, CER1, FOXA2, IRX3* and *NKX2-1* along the time course (**Fig. 2f**). Importantly, genes that are characteristic to hepatoblasts and hepatocytes, including *GATA6, AFP, FGB* and *APOB* were markedly up-regulated, some already during the second stage of the protocol (days 7-10) as demonstrated by *APOB* (**Fig. 2e, f**). These were apparent in Louvain clusters without expression of lung markers, namely clusters 0 and 12 (**Suppl. Fig. 2e, f, Supplementary Table 3**). Finally, we performed gene category enrichment analysis to detect candidate pathways underlying the differentiation of lung progenitors. It showed association of TGF-*β*/Smad and Wnt/*β*-catenin with the formation of DE, the Hippo pathway with foregut differentiation, and Notch signaling in conjunction to insulin-like growth factor signaling with the NKX2-1+ lung progenitor cells (**Fig. 2g**). Taken together, these results indicated that the specification of hepatoblasts originate at an earlier stage than foregut formation and lung progenitors without requiring stimulation by exogenous FGF2, BMP4, or HGF (Carpentier et al., 2014; Hannan et al., 2013).

### Early transcriptional regulators of lung and hepatoblast specification

We next sought to identify genes and mechanisms that promote the emergence of lung progenitors, and the formation of liver cells at the DE stage, by focusing on the timing of up-regulation of known TFs. In this regard, *Isl1* has recently been characterized as a key regulator of *Nkx2-1* in the early foregut (Kim et al., 2019), and *Hhex* has been associated with the activation of *Hnf4* TFs which are essential for liver development (Hunter et al., 2007). Moreover, the TFs *Gata6, Foxa1* and *Foxa2* are well known for their crucial roles in the formation of the liver (Li et al., 2009; Zhao et al., 2005). Analysis of scRNA-seq data of the first 6 days of differentiation revealed the up-regulation of *GATA6, FOXA2* and *HHEX*, but not of *ISL1* or *NKX2-1* (**Fig. 3a,b**). Interestingly, the single cell data showed that only very few cells expressed *T (Brachyury)* in the first 6 days of differentiation (**Fig. 3b**). Because the transient expression of *Brachyury*, followed by *Foxa2*, is known to take place in a subpopulation of the DE called mesendoderm (Burtscher and Lickert, 2009), the absence of *T* indicated that the origin of the liver and lung cells is not associated with mesendoderm origin. Collectively, this indicated that the regulation that promotes the differentiation of liver cells is being established during the early specification of DE rather than upon foregut formation.

**Figure 3:**
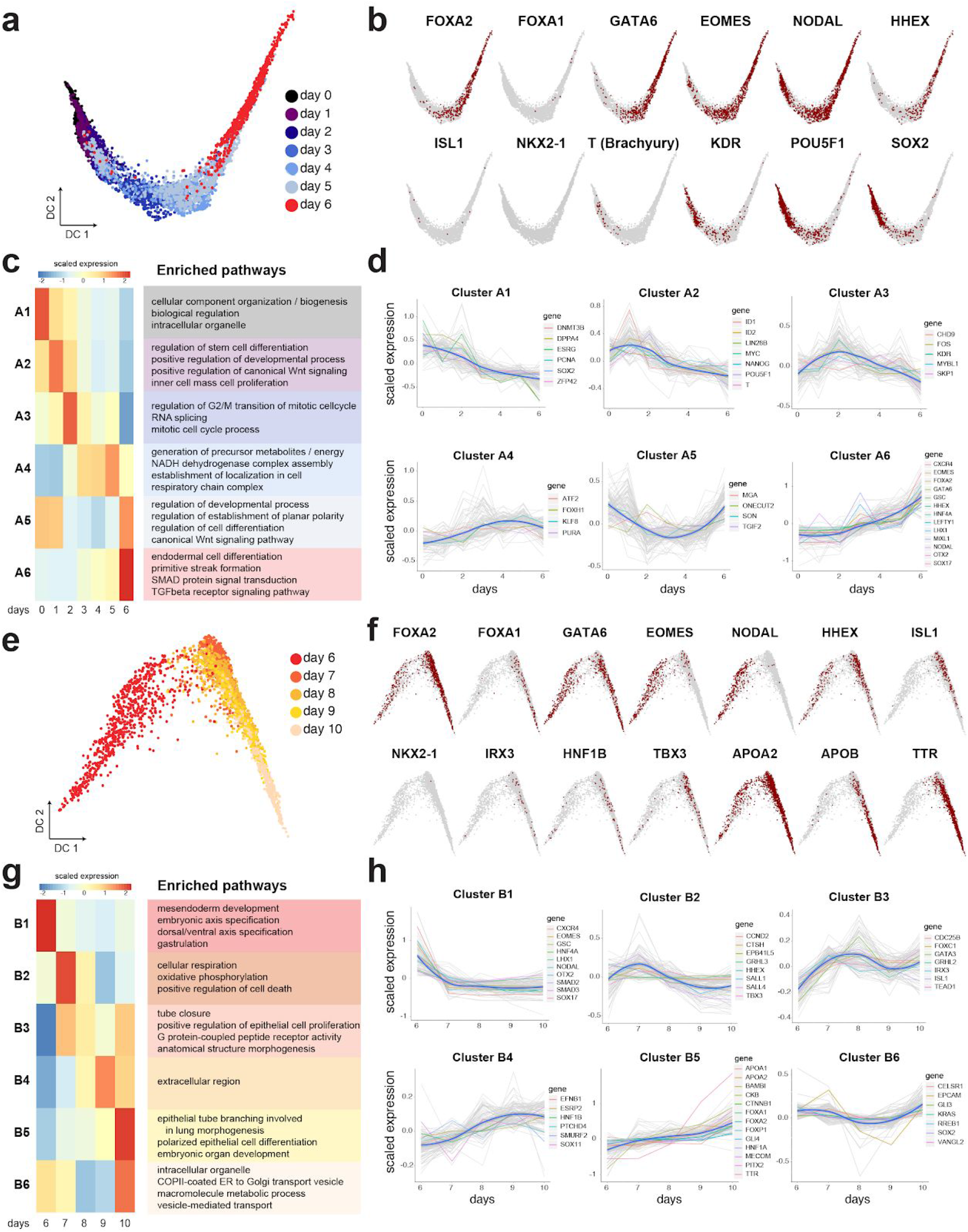
Dissection of the kinetic patterns of genes underlying the differentiation of lung progenitors and hepatoblasts. (**a,b)** Diffusion map of single cells for days 0-6 showing the transition from pluripotency towards DE **(a)**, marked with positive cells for indicated gene markers **(b). (c)** Scaled mean expression per gene cluster and associated pathways identified by hierarchical clustering of the genes with significant associations to days 0-6 (adjusted p value < 0.005, 6101 genes; high-to-low association colored red to blue). (**d**) Scaled mean expression of each gene cluster at the indicated time points. Grey lines correspond to the scaled expression values of the top 100 genes per cluster, and the blue lines correspond to the median expression within the cluster. **(e-h)** Diffusion map and cluster analysis as above for days 6-10 of differentiation (adjusted p value < 0.005, 9352 genes). Genes discussed in the text are highlighted.

To investigate further the circuitry of TFs that promotes DE specification, we used hierarchical clustering of all the genes that exhibited transcriptional changes during the first 6 days of differentiation. We generated 6 major clusters numbered A1-A6, representing different gene expression kinetics, and analyzed the enrichment of respective GO pathways in these clusters (**Fig. 3c,d; Supplementary Table 5**). Expression of major DE genes, such as *EOMES, LHX1, OTX2, CXCR4, LEFTY1, SOX17* followed a very similar pattern with gradual upregulation peaking at day 6 (cluster A6). This cluster was enriched in pathways involved in endoderm differentiation and regulation of SMADs and TGF-*β* signaling (**Fig. 3c, Supplementary Table 5**), including the loop of EOMES-Activin/Nodal signaling (**Suppl. Fig. 3a**). Several studies that have previously analyzed bulk DE cells have suggested that a subset of KDR+ (VEGF receptor 2) progenitors generates hepatic cells from differentiated human ESCs, and that KDR signaling is important for their specification (Gordillo et al., 2015). Contrarily, we have noted that the up-regulation of *FOXA2, GATA4* and *GATA6* coincided with a downward trend of KDR (**Fig. 3b** and cluster A3 in **d**), suggesting that KDR is not involved in instructing the early liver phenotype alongside lung cells in our case.

To next investigate whether lung progenitors emerged during the second stage of the protocol, and whether this process coincided with maturation of hepatoblasts, we used hierarchical clustering to generate 6 clusters of gene expression, numbered B1-B6, for days 6-10 of differentiation (**Fig. 3e-h and Supplementary Table 4**). We noted that the genes linked to the induction of DE, namely *EOMES, LHX1, GSC, OTX2, SOX17* and *CXCR4* (Costello et al., 2015), and GO pathways associated with meso-endoderm development were sharply down-regulated upon dual-Smad inhibition in this stage (cluster B1, **Fig. 3g,h**). An important exception was *FOXA2*, whose expression is essential for development of the foregut. *FOXA2* belonged to cluster B5 together with *FOXP1* and *PITX2* (**Fig. 3h**), which are crucial for lung morphogenesis and asymmetry in the mouse, but not for the initial specification of the lung in the mouse (Lin et al., 1999; Shu et al., 2007). In accordance, GO pathways showed enrichment of lung morphogenesis, embryonic organ development and epithelial cell differentiation in cluster B5 (**Fig. 3g**), even though at this stage NKX2-1 hasn’t been up-regulated yet (**Fig. 3f**). Nevertheless, approximately halfway through this stage, *ISL1* was up-regulated in cluster B3 alongside *IRX3. Isl1* has recently been shown to regulate the development of lung lobes and trachea-esophagus tube separation by the activation of *Nkx2-1* (Kim et al., 2019), and *Irx3* is necessary for lung formation by promoting the proliferation of branched epithelium (van Tuyl et al., 2006). In accordance, the GO patterns of cluster B3 included epithelial cell proliferation and tube closure, which collectively indicated that the lung progenitor program has been initiated between days 6-10 (**Fig. 3g**). *ISL1* is also known to regulate SHH in the foregut endoderm (Lin et al., 2006), which suggested that basal differentiation of NKX2-1-eGFP+ cells was supported by endogenous production of SHH (**Fig. 1b**). The connection to the activation of the SHH pathway at days 6-10 was apparent in clusters with upward trend, including *GLI4* in B5 and *GLI3* in B6 (**Fig. 3h**). Furthermore, comparison of our data to scRNA-seq of the mouse embryo at the time of gastrulation (Pijuan-Sala et al., 2019), showed that *Isl1* and *Irx3* were specifically enriched in the foregut endoderm and the pharyngeal endoderm around E8.0-E.5 (**Suppl. Fig. 3b-d**). In agreement with the foregut association, pancreatic genes such as *PDX1*, which is associated with the midgut and are suppressed by SHH (Deutsch et al., 2001), was not apparent in our data (**Suppl. Fig. 3b-d**).

Lastly, we analyzed hepatoblast and hepatocyte genes in days 6-10 following the activation of *HHEX* in days 0-6. At this stage we noted the upregulation of pioneering liver TFs including, *HNF1A, HNF1B*, and *TBX3* which governs the expansion of the liver bud (Lüdtke et al., 2009), as well as first indications of functional liver genes, namely, *APOB, APOA2*, and Transthyretin (*TTR*) which is secreted by the liver (**Fig. 3f,h**). Importantly, we noted that the expression of these liver genes was associated with Louvain clusters 0 and 12, which were to a large extent mutually exclusive from the above mentioned lung factors and related Louvain cluster 10 (**Fig. 2c and Suppl. Fig. 2f, Supplementary Table 3**). Taken together, these scRNA-seq data indicate that the lung and liver lineages are already set in their respective separate trajectories during the second stage of the protocol.

### Trajectory analysis reconstructs lung and liver branching

We next investigated mechanisms that drive the separation of lung progenitors from the liver fate. Trajectory analysis of differentiation days 11-15 combined with days 7-10 revealed a branching event in the high dimensional single cell data manifold (**Fig 4a**,**b and Suppl. Fig. 4**). The lung progenitors appeared to become separated at the intersection of the FE stage with the last stage, and the maturity of lung and liver fates increased progressively over time when compared to NKX2-1eGFP+ and NKX2-1eGFP-bulk populations **(Fig. 4c, d**). Importantly, and as discussed above, the hepatoblast fate preceded the lung identity and was already established at days 7-10.

**Figure 4.**
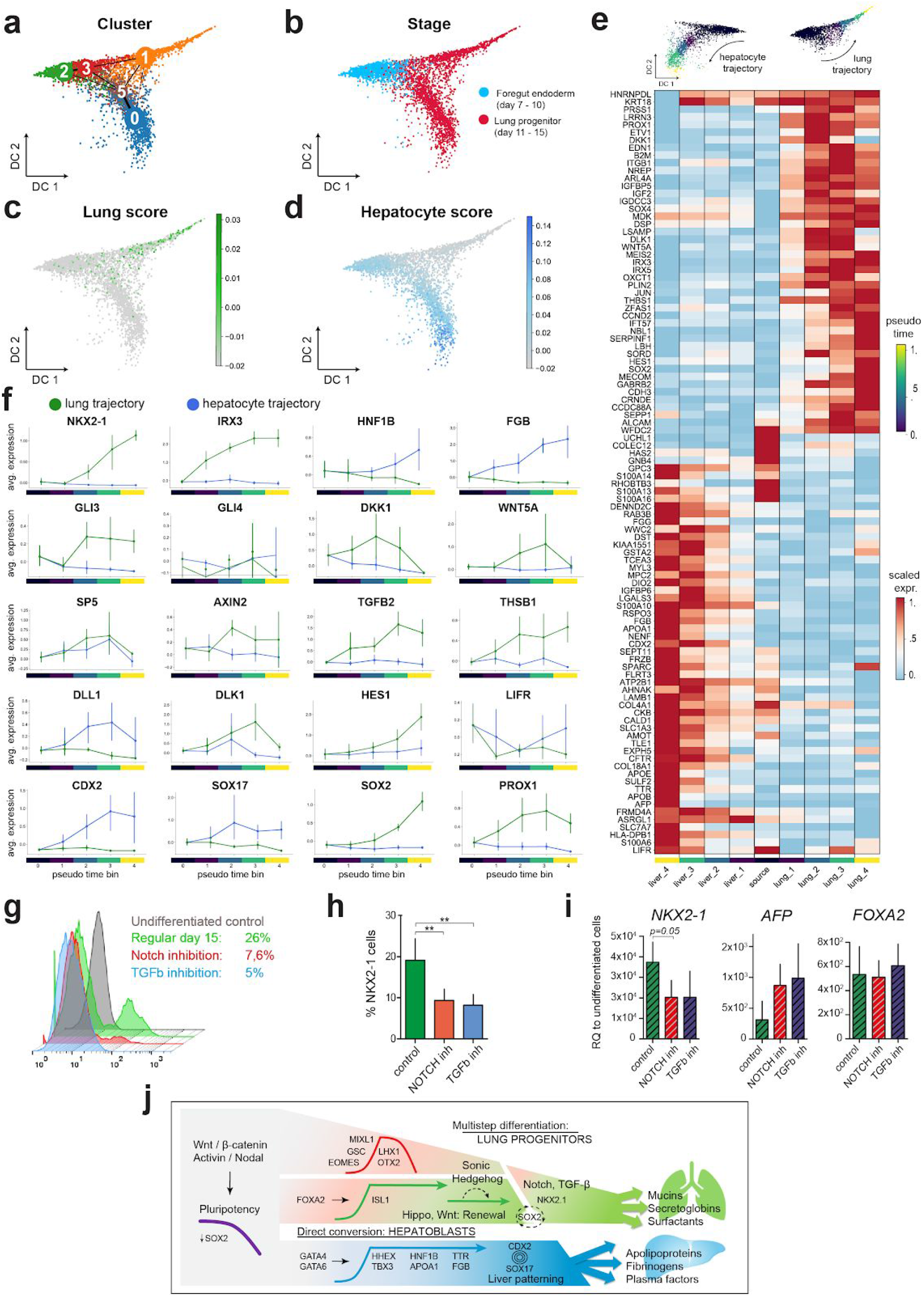
Reconstructing the transcriptional transitions from pluripotency to lung progenitors and hepatoblasts. **(a, b)** Diffusion maps and Louvain clustering representing single cell transcriptomes of days 7-10 (blue) and 11-15 (red) in **(b). (c, d)** Color coded scores indicate similarity of single cell transcriptomes to bulk mRNA-seq data of NKX2-1-eGFP+ (**c**) and eGFP-populations (**d**). **(e)** A heatmap showing unsupervised clustering of top differentially expressed genes along the lung and liver pseudotime trajectory respectively. **(f)** Line plots showing the expression dynamics of the indicated genes in the respective branches in accordance to the binned pseudotime ordering (vertical lines represent confidence intervals of 95%). **(g**,**h)** Representative flow cytometry and the quantification of NKX2-1-eGFP+ lung progenitors on day 15 of differentiation, with or without the treatment by DAPT or SB431542 as indicated in days 11-15 (bars represent mean ± SD, n = 4 biological replicates, **p ≤ 0.01 by unpaired, one-tailed t-test. **(i)** The respective fold change of *NKX2-1, AFP* and *FOXA2* on day 15 relative to the parental undifferentiated cells (quantified by qRT-PCR 2^(–ΔΔCT)^, bars represent mean ± SD, n = 3 biological replicates, unpaired, one-tailed t-test. **(j)** A graphical summary of genes, their phases of expression, and mechanisms underlying the differentiation of lung progenitors and hepatoblasts from pluripotency state.

The pseudotime trajectories in the last stage of the differentiation protocol revealed key differences in the lung and liver differentiation branches (**Fig. 4e, f**). As expected, the expression of lineage specific markers such as *NKX2-1, IRX3*, and *HNF1B*, and *FGB* was exclusively increased along the two trajectories (**Supplementary Table 5**). We noted that *GLI3*, a key transcriptional repressor that is regulated by SHH signaling, was upregulated only in the lung branch. This corresponded to the exogenous and endogenous activation of the pathway by SHH and *ISL1* (**Fig. 3g,h**). Moreover, key components of the Wnt/*β*-catenin pathway, including *DKK1, WNT5A, SP5*, and notably the pathway’s canonical marker *AXIN2*, exhibited considerably higher expression in the lung trajectory (**Fig. 4f**). The specificity of these pathways to the lung trajectory is particularly interesting given that the exogenous treatment by SHH and CHIR (**Fig. 1a**) did not activate the pathways in the neighboring hepatoblasts (**Fig. 1j**). Considered together with earlier studies that revealed central roles of SHH and Wnt/*β*-catenin in foregut-early lung development (Goss et al., 2009; Harris-Johnson et al., 2009; Motoyama et al., 1998), these results indicate that early hepatocytes might be refractory to these pathways. In accordance, the exclusive expression of *SOX2* in the lung trajectory is another indication of the lineage specific activity of Wnt/*β*-catenin because SOX2, canonical Wnt signaling, and FGFs often intersect in the regulation of self-renewal in development (Turner et al., 2014) (**Fig. 4f**).

Next, we found evidence for the activation of the Notch pathway specifically in the lung branch. This was inferred by us through the lung branch specific expression of *HES1*, a canonical TF of the Notch pathway, as well as the Notch pathway modulator *DLK1* that is known to be involved in lung branching and morphogenesis (Falix et al., 2012). To mechanistically test if indeed Notch is needed for lung specification in this system, we inhibited the Notch pathway by the γ-secretase inhibitor DAPT from day 11 onwards (**Fig. 4g-i**). In accordance with our hypothesis, this led to a significant decrease in the number of NKX2-1eGFP+ progenitors, a reduction in *NKX2-1* expression, and a reciprocal increase of *AFP* expression. Interestingly, a key ligand of the Notch pathway, namely *DLL1*, was specifically expressed in the liver branch (**Fig. 4f**), which could indicate that hepatoblasts promote Notch signaling in lung progenitors by paracrine signaling. Moreover, we inhibited the TGF-*β* pathway using SB431542 after noting that the expression of *TGFB2* and *THSB1* was specific to the lung branch (**Fig. 4f**). We observed a similar effect as with the Notch pathway inhibition, namely, a decline in the number of NKX2-1-eGFP+ progenitors and expression of *NKX2-1* in treated cells (**Fig. 4g-i**). Taken together, this proposes an important role for a crosstalk between Notch and TGF-*β* in the specification of the foregut towards lung progenitors.

Finally, the pseudotime trajectory ordering has indicated surprising involvement of key developmental genes that to our knowledge were previously not implicated in the specification of hepatoblasts and lung progenitors. We noted that *CDX2, SOX17*, and LIF/STAT3 signaling via *LIFR* expression were highly enriched specifically in the liver trajectory (**Fig. 4f**), while mouse knockout studies have not made observations pertaining to the primary roles of these factors in liver development (Kumar et al., 2019; Onishi and Zandstra, 2015; Spence et al., 2009). Moreover, the expression of *PROX1* which is considered an early and specific marker for the developing liver and pancreas in the mouse foregut (Burke and Oliver, 2002), was in fact specifically expressed in the lung trajectory (**Fig. 4f**). In addition, the expression of *FOXP1* and *PITX2* included both the lung and liver trajectories, and *FOXP2* was not detected (**Suppl. Fig. 4e**), contrary to the specific expression in lung progenitors described in the mouse embryo (Lin et al., 1999; Shu et al., 2007). This indicated differences in the regulation of key developmental genes between mouse and human, which we incorporated in a comprehensive roadmap model for human lung and hepatoblast differentiation from the pluripotency state (**Fig. 4j**).

## DISCUSSION

Lung disease is one of the leading causes of death. Lung organoid model systems based on use of PSCs promise to greatly increase our ability to study lung disease. Likewise, harnessing the mechanisms of lung development will advance tissue engineering and regenerative medicine if we understand the fundamental differentiation processes during the formation of the lung and upon injury. Time-series single cell RNA-seq has been recently used to model the differentiation of human lung progenitors towards alveolar cell identities (Hurley et al., 2020). In this work we map even earlier molecular processes in human lung development by reconstructing genes, networks, and pathways, in the transition from pluripotency to the NKX2-1 lung progenitor state.

Of key interest in the formation of lung progenitors is how the expression of *NKX2-1* is initiated in the foregut. Our analysis points to activity of FOXA2 and ISL1 in this process. We find that *FOXA2* is an epiblast-stage TF whose expression persists after the inhibition of TGF-*β* signaling in the second stage of the protocol (leading to abrupt down-regulation of *EOMES, MIXL1, GSC*). Interestingly, *FOXA2* has been shown to directly activate *ISL1* in cardiac progenitors (Kang et al., 2009), and *ISL1* was shown to activate *NKX2-1* in the early foregut in a process that is required for lung formation (Kim et al., 2019). Moreover, Shh signaling is required for the development of the foregut (Litingtung et al., 1998; Motoyama et al., 1998), ISL1 has been shown to regulate *Shh* expression in the foregut (Lin et al., 2006), and here we show that SHH treatment increases the number of NKX2-1 progenitors. The connection between ISL1 and SHH signaling is substantiated also by the similar developmental defects of the cardiovascular system in *Isl1* and *Smo* (*Smoothened*) mutant mouse embryos (Lin et al., 2006). Importantly, complexes of Foxa2, Otx2 and Lhx1 have been implicated in autoregulation and pioneering the transcription of foregut genes (Costello et al., 2015). Based on the similar expression kinetics of *FOXA2* and *OTX2* and the up-regulation of *LHX1* observed in our data, we reason that a similar process might control the progenitor differentiation of lungs *in vitro*. This is illustrated in our model: it places FOXA2 and OTX2 in an autoregulatory loop during the formation of early DE which induces the expression of ISL1 and subsequently NKX2-1. ISL1 in parallel up-regulates and activates SHH signaling which *in vivo* has been shown to promote lung morphogenesis and branching patterns (Litingtung et al., 1998) (**Fig. 4J**).

The emergence of both lung and liver cells in the differentiation protocol posed the question if a common progenitor can be identified. Based on our trajectory analysis we found no evidence indicating the formation of bi-potent lung-liver progenitors. We found that the inception of hepatoblast transcription networks takes place much earlier than the initiation of the lung program. The induction of *HHEX, HNF4A* and *HNF1B* are unmistakable signs of hepatoblast differentiation, and remarkably these genes were up-regulated as early as 3-4 days of DE differentiation, while the earliest signs of *ISL1* upregulation appeared only 4-5 days later. Therefore a mechanism whereby a common progenitor population responds to cues that promote a choice between alternate fates, namely of lung or liver, is unlikely here.

Our data indicates a direct conversion of early DE to hepatoblasts. Mechanistic mouse studies show that the activation of the Activin / Nodal – EOMES loop turns on the high levels of canonical TFs that control hepatogenesis in the mouse embryo. This includes *Gata6* and *Gata4* (Zhao et al., 2005), *Foxa2* and *Foxa1* (Lee et al., 2005), which have been implicated in pioneering chromatin remodeling in foregut precursors (Zaret and Carroll, 2011), and were utilized in the induction of hepatocyte transdifferentiation (Chen et al., 2018; Wamaitha et al., 2015). Moreover, *Gata4* and *Gata6* were shown to be functionally redundant in promoting the development of the liver, and together with *Foxa2* they promote liver specification by inducing *Hhex* (Denson et al., 2000). Remarkably, in our data a number of hepatoblast TFs e.g. *HNF4A, TBX3*, and liver factors that are secreted into the plasma e.g. *APOA2, FGB, TTR, APOB* were induced shortly after *HHEX*, which is evidence that may support our “fast track” liver differentiation model from early DE (**Fig. 4J**). Interestingly, the expression of *HHEX* itself ceased almost entirely in the third phase of the protocol, despite the continued expression of *FOXA2* and *GATA6*. It is possible that a majority of cells have undergone specification already at the beginning of the third phase of the protocol, or that the disappearance of *HHEX* is connected to the downregulation of *OTX2*, which directly regulates the *HHEX* promoter foregut cells (Rankin et al., 2011), or that a different mechanism is at play.

Of special interest for tissue engineering and disease modeling is the association of the *FOXA2* -*ISL1* -*NKX2-1* trajectory with developmental pathways that regulate patterning and self-renewal in the lung (**Fig. 4J**). Our evidence of i) the up-regulation of *SOX2* in the lung branch that coincided with activation of Notch and Wnt/*β*-catenin signaling, ii) activation of the Hippo pathway during the foregut stage of the differentiation protocol, iii) the reduction of NKX2-1 progenitors upon TGF-*β* and Notch inhibition, together with iv) Yap dependent TGF-*β* mediated induction of proliferation by Sox2 in mouse airway progenitors (Mahoney et al., 2014), create a putative picture of renewal mechanisms that might inform the future derivation of human lung stem cells. Moreover, the specific expression of *SOX17* and *CDX2* in the liver trajectory is novel and important. In the mouse *CDX2* expression is restricted to the hindgut where it serves to regulate intestinal stem cell development (Kumar et al., 2019). Therefore, the up-regulation of *CDX2* in conjunction to *SOX17*, which serves as a patterning factor in the liver bud (Spence et al., 2009), might indicate that these factors have central roles in the regulation of human liver development. Comparing the trajectories of additional foregut-lung TFs with their associated functions in mouse development identified additional differences. Specifically, we noted the broadening of the expression profiles of *FOXP2* and *PITX2*, which might imply that evolutionary changes increased their involvement in the differentiation of the human foregut. Another example is the association of *PROX1* with lung progenitors as opposed to hepatocytes in the mouse (Gordillo et al., 2015). This reveals putative mechanisms that are specific for the formation of human foregut precursors and their differentiation to lung and liver progenitors.

In summary, the high temporal resolution of our single cell trajectory analysis enabled us to construct a hierarchical model of gene expression changes along two trajectories from hPSC to lung and liver fate respectively. In our analysis the pseudotime estimation agreed well with the highly resolved time series of sampling. Inferring our results in light of *in vivo* studies provided a detailed understanding of mechanisms, their timing, and evolutionary changes that regulate the specification of lung progenitors in the foregut. These findings should inform applications of iPSCs in tissue engineering efforts for regenerative medicine of the lung, possibly including the derivation of lung stem cells. Moreover, our work provides a high resolution “roadmap” for the direct conversion of hepatoblasts, which could be used for improving the manufacturing of hepatocyte grafts in liver disease.

## Supporting information

Table S1 List and catalogue numbers of reagents

Table S2 Single-cell level meta data

Table S3 Marker genes of louvain clusters

Table S4 Results of spline fit

Table S5 Hierarchical clustering of genes

Table S6 Differential gene expression along trajectories

Table S7 Gene-Gene correlations

## Acknowledgements

We are grateful for the iPSC line provided by Dr. Ruth Olmer and Dr. Prof. Ulrich Martin (Hannover Medical School). We would like to thank Lukas Simon and Malte Lücken (Helmholtz Center Munich) for helpful advice on statistical modelling and integration across stages, as well as all the members of the Theis lab for valuable input and discussion regarding the analysis of single cell data. This work was supported by the Helmholtz Association and the German Center for Lung Research (DZL).

## Data and Code availability

The scRNA-seq data used in this study has been deposited in the Gene Expression Omnibus database under accession number GSE167011.

Code to reproduce the analyses and figures generated in this manuscript can be assessed via https://github.com/theislab/2020_iPS_Lung_Differentiation.

## Author contributions

M.D. and H.B.S. conceived and supervised the entire study. C.O., M.D. and H.B.S. wrote the manuscript. C.O. performed the majority of experiments. I.A. generated single cell RNA-seq data. M.A. performed analysis of single cell data. H.B.S. and F.J.T. supervised the analysis of single cell data.

## SUPPLEMENTARY FIGURES

**Supplementary Figure 1:**
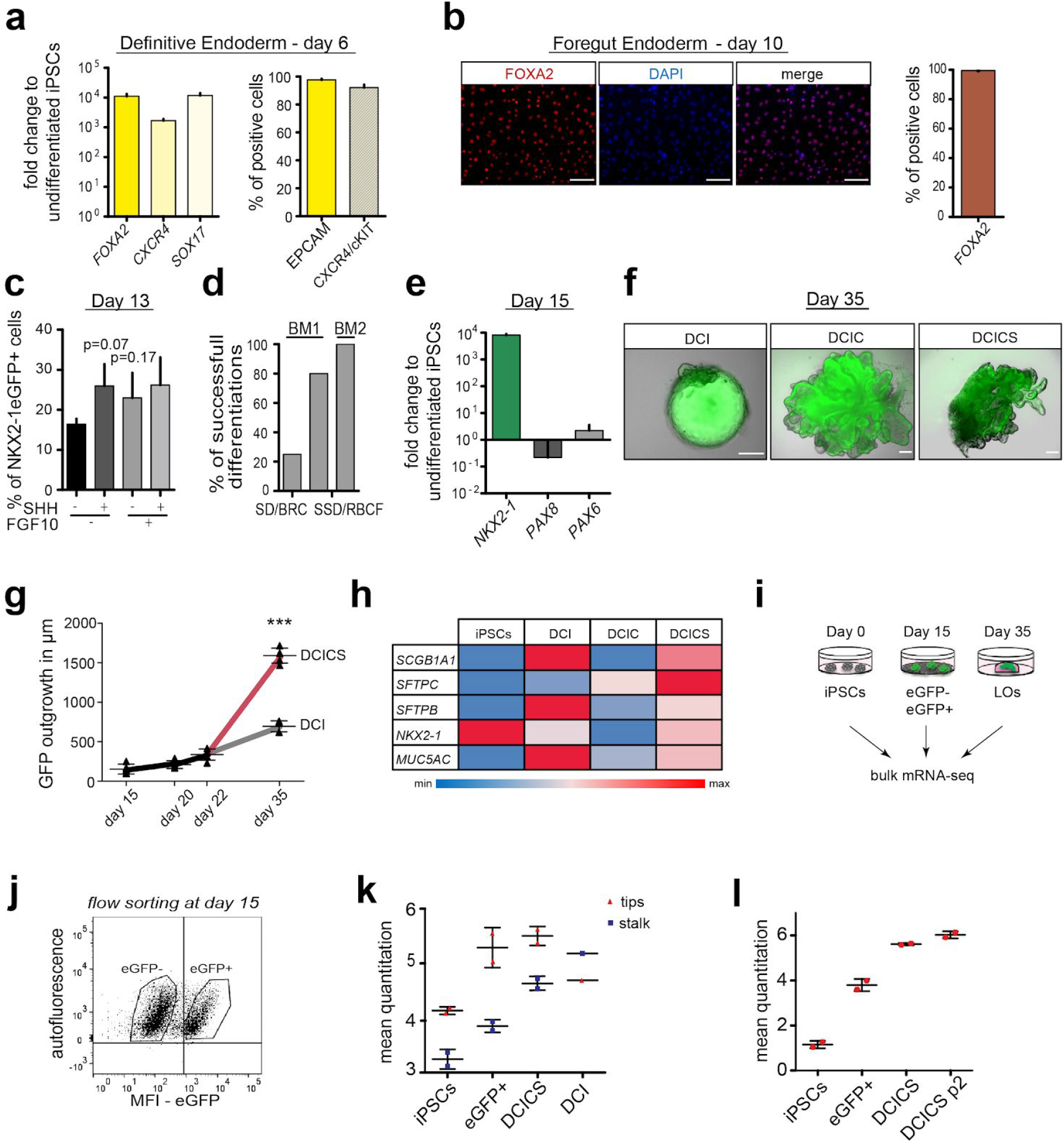
Related to Fig. 1. **(a)** The differentiation of DE precursor cells analyzed by qRT-PCR and flow cytometry of cell surface markers on day 6 of differentiation (undifferentiated parental cells were used for normalization and gate settings). **(b)** Representative microscopic images and quantification of foregut precursors analyzed by immunostaining of FOXA2 and DNA staining (DAPI in blue) on day 10 of differentiation (scale bars: 100 µm). **(c)** Quantification of NKX2-1-eGFP+ cells analyzed on day 13 of differentiation by flow cytometry; treatment by SHH and FGF10 as indicated (n=3 biological replicates in **a-c**). **(d)** Percentage of successful differentiation experiments with the indicated conditions corresponding to **Fig. 1b. (e)** Analysis of markers representing lung, thyroid and forebrain tissues *NKX2-1, PAX8* and *PAX6* respectively (normalized and analyzed as in **a**; n = 4 biological replicates). **(f-h)** Fluorescent microscopy images (**f**) and organoid growth curves **(g)**, and qRT-PCR **(h)** of markers characteristic of the early lung. Organoids were created from day 15 colonies using conditions outlined in **Fig. 1a** of NKX2-1-eGFP+ cells, and were analyzed on day 35 (scale bars: 200 µm in **f**; qRT-PCR normalized as in **a**, n = 5 biological replicates; ***p ≤ 0.0001 unpaired, one-tailed t-test). **(i)** Schematic illustration of samples prepared for bulk mRNA-sequencing, and **(j)** representative flow cytometry plots with the gating strategy used for sorting eGFP+ and eGFP-cells (undifferentiated parental cells were used as negative controls to set gates). **(k)** A plot displaying the correlation of the mean quantitation for each data store to tip (red) and stalk (blue) gene signatures from (Nikolić et al., 2017), and **(l)** to the panel of early lung genes shown in **Fig. 1F**. As indicated by an increase of a cohort of fetal lung and alveolar genes, passaging of DCICS-organoids (indicated by p2) further promoted their maturation. Error bars represent mean ± SD in all panels.

**Supplementary Figure 2.**
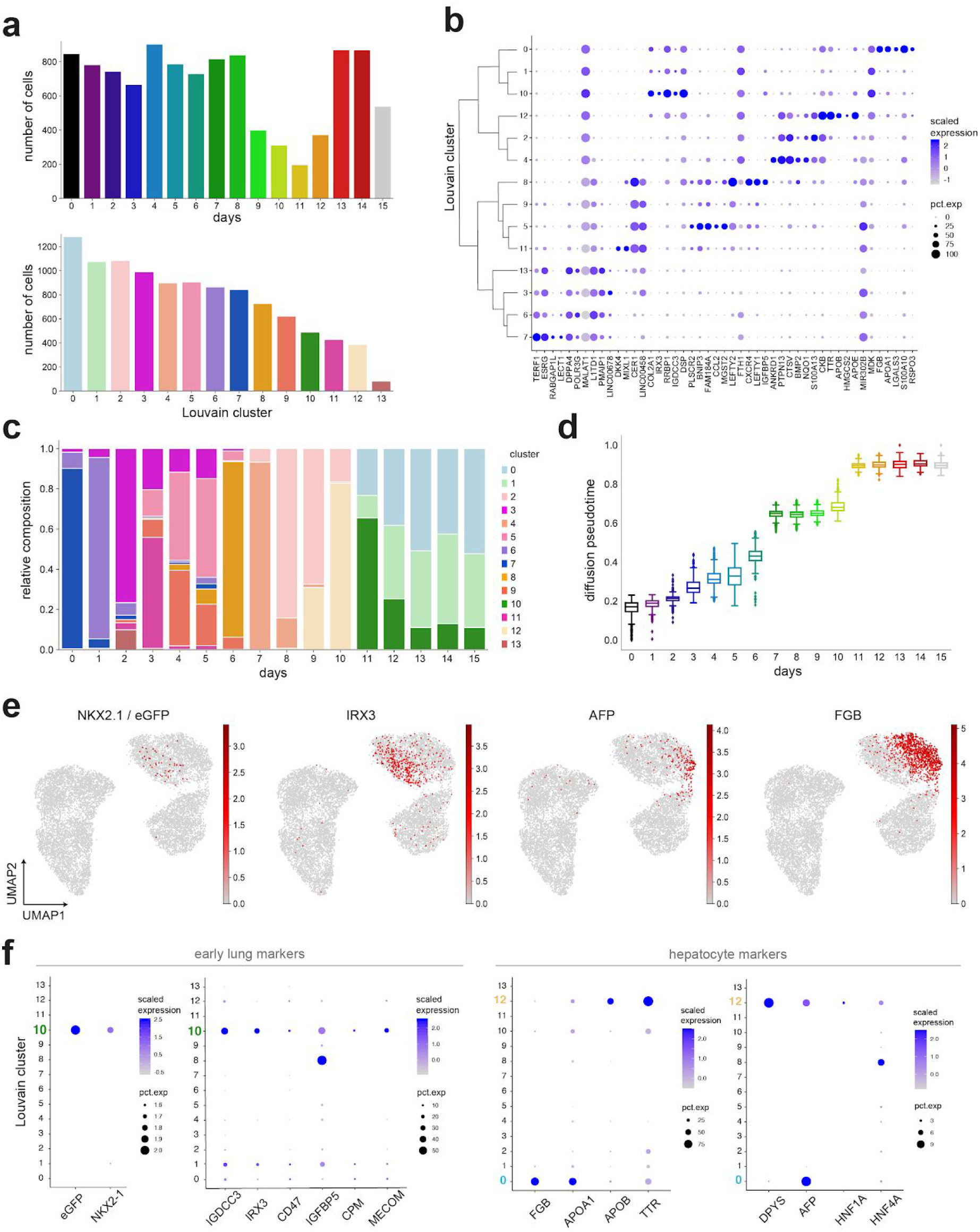
Related to Fig. 2, Overview of the single cell dataset after filtering. **(a)** Barplots displaying the number of cells included in downstream analysis after quality control filtering (details in the experimental procedures) per time point (top), and Louvain cluster (bottom). **(b)** Top 5 genes per cluster shown in a dendrogram created by unsupervised clustering displaying the normalized expression level (across clusters in blue) and the percentage of cells expressing the respective gene according to dot size. **(c)** A barplot showing the Louvain cluster composition according to time point (day) of sampling. **(d)** A boxplot of diffusion in the pseudotime of all sampling time points with boxes representing the interquartile range, horizontal line the median, and the whiskers representing 1.5 times the interquartile range **(e)** UMAPs overlaid with cells expressing lung progenitor, *NKX2-1* and *IRX3*, and liver markers AFP and *FGB*, showing the separation of the two lineages. **(f)** A dot plot indicating that markers of the early lung show higher expression levels in Louvain clusters 10 and 1 than markers of hepatocytes 0 and 12.

**Supplementary Figure 3.**
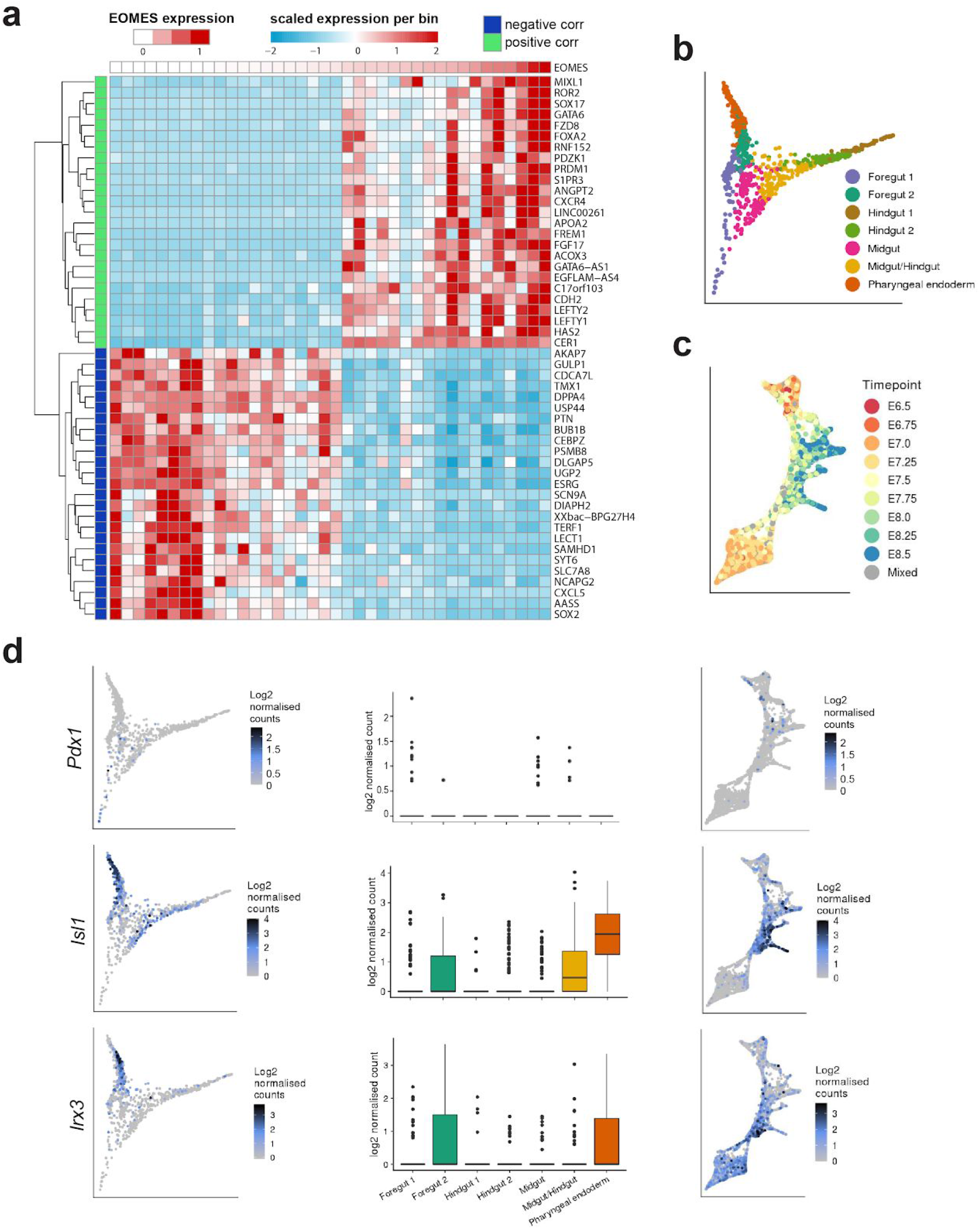
Related to Fig. 3. **(a)** A heatmap encompassing top 25 correlated and anti-correlated genes to *EOMES* expression levels for the respective days 0 to day 6. Cells were clustered into 38 bins based on transcriptomic profile, scaled average expression per bin is shown ordered according to *EOMES* expression. This revealed an association of *EOMES* with *LEFTY1, LEFTY2, CER1*, and *ROR2*, indicating active EOMES-Activin/Nodal circuit (Teo et al., 2011). **(b-d)** Analysis that is based on scRNA seq database of the mouse embryo during organogenesis (Pijuan-Sala et al., 2019), including **(c)** representative time points of embryo collection, and the **(d)** gene expression levels and tissue association of *Pdx1, Isl1* and *Irx3*.

**Supplementary Figure 4.**
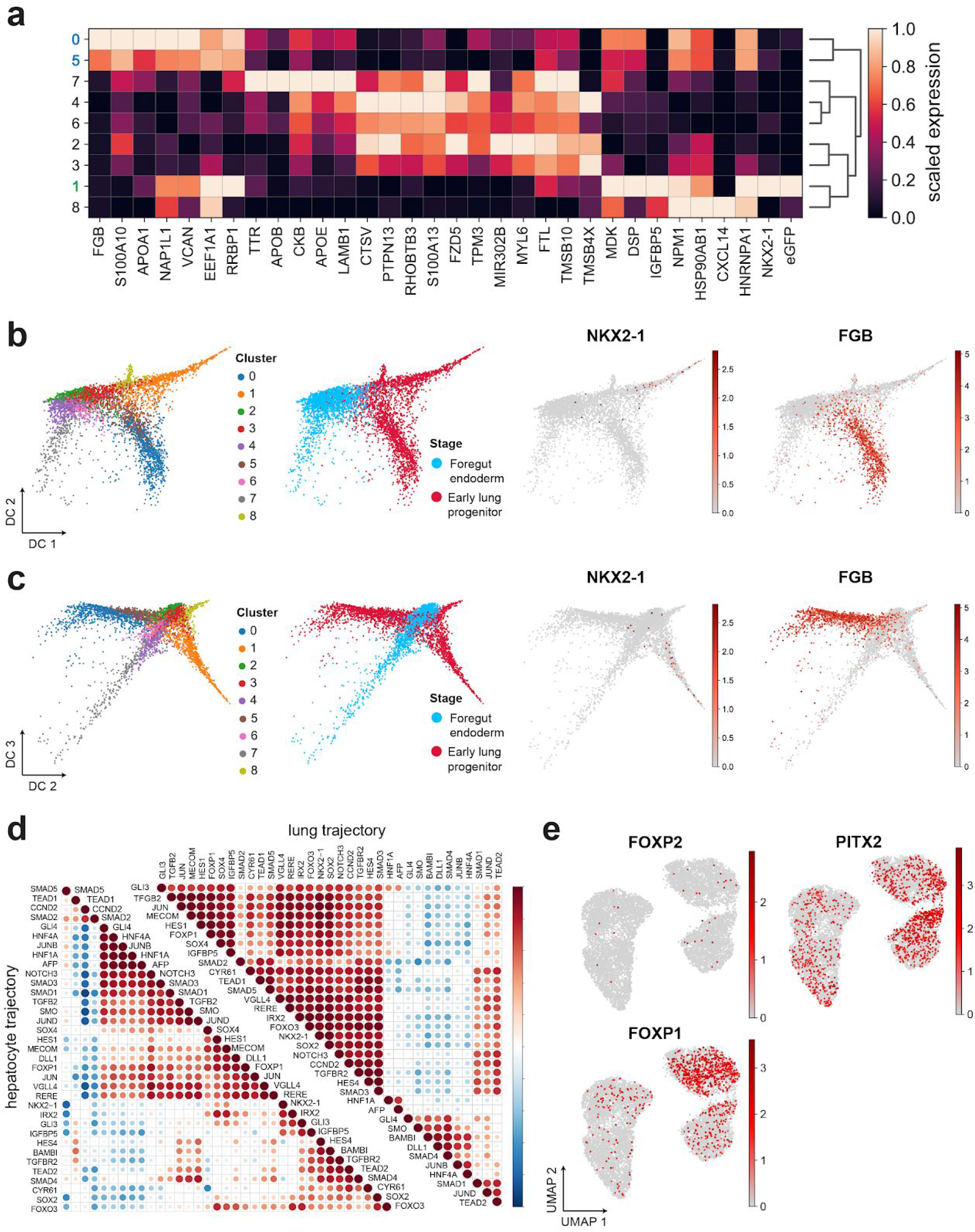
Diffusion map of all cells from days 7-15 of differentiation reveals minor branches that were not included in Fig 4. **(a)** Unsupervised clustering of the top 5% DE genes in 9 Louvain clusters. **(b**,**c)** Diffusion maps showing that the lung marker *NKX2-1* (and *NKX2-1-eGFP*) was restricted to sub branch #1, and hepatocyte marker *FGB* was enriched in the branches encompassing clusters 0 and 5. Cluster colors in **(a)** correspond to lung (green) and liver (blue) branches in **Fig. 4**. Diffusion map with diffusion component 1 vs. 2, and 2 vs. 3 are shown in **(b)** and **(c)** respectively of cells from day 7 to 15 coloured by louvain cluster (left) and stage (middle-left). **(d)** Pairwise gene pearson correlation profiles along the lung and liver pseudotime trajectories. **(e)** UMAP of all time points indicating expression levels of FOXP2, PITX2 and FOXP1.

## EXPERIMENTAL PROCEDURES

### Stem cell maintenance and passaging

Human iPSC line *NKX2-1*^*eGFP/+*^ was kindly provided from Hannover Medical School (Olmer et al., 2019), maintained in feeder-free conditions, in StemMACS iPS-Brew XF (Miltenyi Biotech) and passaged with Accutase (Sigma-Aldrich) on tissue cell culture plates pre-coated with 1:100 dilution of Geltrex Basement Membrane Matrix in DMEM/F-12 (both from ThermoFisher Scientific).

### Differentiation of hiPSCs into NKX2-1+ early lung progenitors

To induce definitive endoderm differentiation, hiPSCs were maintained in iPSC-Brew and when reached 80% confluency (day 0), the cells were rinsed with DPBS and incubated in Accutase (Sigma-Aldrich) for 10 minutes, at 37°C. The detached cells were triturated into single-cell suspensions and seeded onto 24-well plates pre-coated with 1:40 Growth Factor Reduced (GFR) Matrigel (Corning), in a density of 2 × 10^5^ cells/cm^2^. Cells were immediately treated with 100 ng/ml activin-A, 1 μM CHIR99021, and 10 μM Y-27632 (all from R&D Systems), in Definitive Endoderm Basal Media (DE-BM) consisting of RPMI1640 medium, 1 x B27 supplement, 1 x NEAA and GlutaMAX (all from ThermoFisher Scientific). On days 1-6 the DE-BM was supplemented with 100 ng/ml activin-A, 1 μM CHIR99021 (R&D Systems) and 0.25mM (day 1) and 0.125 mM (days 2-6) sodium butyrate (Sigma-Aldrich).

For the foregut endoderm stage the Basal Mediums (FE-BM1 and FE-BM2) tested, were prepared as follows, FE-BM1: DMEM/F12, 1 x GlutaMAX, 1 x B-27 and N-2 supplements, 50 U/ml of penicillin/streptomycin, 0.05 mg/ml of L-ascorbic acid (Sigma-Aldrich) and 0.4 mM of monothioglycerol (Sigma-Aldrich) (Gotoh et al., 2014) or FE-BM2: 75% IMDM, 25% Ham’s F-12, B27 supplement and N2 supplement, 0.05% bovine serum albumin, 1 x GlutaMAX, 50 U/ml of penicillin/streptomycin (all from ThermoFisher Scientific), and 0.05 mg/ml of L-ascorbic acid, 0.4 mM of 1-thioglycerol (Sigma-Aldrich) (Hawkins et al., 2017) (Fig. 1b). On day 6 DE cells were collected with Accutase for 10 minutes at 37°C and re-plated in a density of 1:2-1:4 onto GFR Matrigel-coated plates. The cells were treated for 2 days (days 6, 7) with 50 ng/ml sonic hedgehog (SHH; R&D Systems), 2 μM dorsomorphin (DSM; Tocris) and 10 μM SB431542 (Miltenyi Biotec), supplemented with 10 μM Y-27632 (R&D Systems) for the first 24 hours. On day 8 the medium was changed to FE-BM1 or 2 respectively, with DSM and SB431542.

To induce the lung progenitor (LP) stage, on the day 10, the medium was switched to FE-BM1 or BM2 containing 20 ng/ml recombinant human BMP4 (rhBMP4, R&D Systems), 50 nM retinoic acid (RA; Sigma-Aldrich), 3 μM CHIR99021 and 20 ng/ml rhFGF10 (R&D Systems). For the inhibition of Notch and TGF-*β* pathways, 10 μM SB431542 and 100 μM DAPT (TOCRIS Bioscience) were additionally used at the LP induction stage of differentiation respectively. Growth media and supplements are listed on Supplementary Table 1a.

### Flow cytometry analysis and sorting of iPSC-derived lung progenitors

The cells were rinsed with DPBS and incubated in Gentle Cell Dissociation medium (STEMCELL Technologies)/0,05%Trypsin (ThermoFisher Scientific) (2vol:1vol) for 10 min, at 37°C. The detached cells were diluted in a FACS buffer containing 2% FBS/DPBS and centrifuged at 200 RCF for 3 minutes, at room temperature (RT). Cell pellets were washed with FACS buffer and centrifuged again. For analysis, the samples were incubated (if necessary) with the conjugated antibodies diluted in FACS buffer, for 25 min on ice, washed twice with FACS buffer and analyzed. Live cells were distinguished by staining with propidium iodide (PI) or Sytox blue (SB). For sorting, the dissociated cells were re-suspended in FACS buffer, stained with PI and sorted into 2% FBS/DMEM. The measurements were obtained using a BD FACSAria II flow cytometer (BD Biosciences). Isotype controls were used for gating stained cells, whereas undifferentiated cells that do not express eGFP were used as negative control for gating NKX2-1^eGFP+^ cells.

### Generation of 3D cultured lung organoids

On day 15 of the 2D differentiation, the cells were rinsed with DPBS and the eGFP+ colonies were dissected with a needle, picked and transferred to 80% Matrigel (diluted in FE-BM2). Drops of 40μL were pipetted into the center of each well of a 24 well tissue culture plate and incubated for 30’ at 37°C. Then, FE-BM2 media supplemented with 10 ng/ml FGF10, 10 ng/ml FGF7 and 3 μM CHIR99021 was added to each well for 7 days. On day 22 the medium was switched to FE-BM2 media supplemented with 50nM dexamethasone (Sigma-Aldrich), 0.1 mM 3-Isobutyl-1-methylxanthine (IBMX; Sigma-Aldrich), 0.1 mM 8-Bromoadenosine 30,50-cyclic monophosphate sodium salt (cAMP; Sigma-Aldrich) and ± 10 μM SB431542, 3 μM CHIR99021. For all the 3D Matrigel culture conditions the medium was changed every second day.

The PneumaCult™ Air Liquid Interface (ALI)-medium (STEMCELL Technologies) was used to induce the bronchial (goblet and/or ciliated cells). In particular, eGFP+ colonies were transferred on day 15 in 1:2 GFR Matrigel/FE-BM2 and placed in 6.5mm inserts, in a total amount of 100 ul. After incubation for 20min in 37°C, FE-BM2 supplemented with 10 ng/ml FGF10, 10 ng/ml FGF7 and 3μM CHIR99021 was added in the lower chamber until day 27. On day 28, the medium was switched to PneumaCult™-ALI Maintenance Medium which was prepared according to the manufacturer’s instructions supplemented with 10μM DAPT (TOCRIS Bioscience) (Firth et al., 2014; Konishi et al., 2016). Growth media and supplements are listed on Supplementary Table 1a.

### Immunostaining

For imaging 2D cultures cells were cultivated on removable 8-well chamber glass slides (Ibidi), rinsed twice with DPBS and fixed with 4% PFA (Sigma-Aldrich)/DPBS for 20 min at RT. Cells were permeabilized with 0.2% Triton X-100 (ThermoFisher Scientific), for 5min and blocked with 3% BSA for 30 min at RT. Then, cells were incubated with primary antibodies diluted in solution containing 0.1% Triton X-100/3% BSA overnight at 4°C, followed by three washes with DPBS, after which they were incubated with secondary antibody in 0.1% Triton X-100/3% BSA solution for 1 hour at RT and washed three times with DPBS. The removable wells were discarded and ProLong Antifade Mountant with DAPI (Thermo Fisher Scientific) was used to mount the coverslip.

For imaging lung organoids, the samples were extracted from the Matrigel drops, upon incubation with 2mg/ml Dispase IV (Sigma-Aldrich), at 37°C for 1h. Then, the organoids were fixed with 4% paraformaldehyde/DPBS for 2 hours at 4°C and incubated in 30% sucrose/DPBS overnight at 4 °C. The samples were embedded in OCT compound (Sakura Finetek) and frozen in -20°C. The frozen samples were cryosectioned into 6-μm slices, permeabilized and blocked with 0.5% Triton X-100/5% Normal Goat or Donkey Serum (NGS or NDS) for 30 min in RT. Next, the slices were incubated with the primary antibodies, diluted in 0.5% Triton X-100/5% NGS or NDS, overnight at 4°C, followed by washing three times with DPBS (10 min each), and stained with secondary antibodies for 3h at RT. After washing twice more, a coverslip was mounted using ProLong Antifade Mountant with DAPI. All antibodies used for the study are listed on Supplementary Table 1b.

### Quantitative RT-PCR

For the RT-qPCR analysis, the cells were lysed and subsequent RNA extraction was performed using the RNeasy Mini Kit (Qiagen) according to manufacturer’s instructions. The RNA was reverse transcribed into cDNA using the Verso cDNA Synthesis Kit (ThermoFisher Scientific) and qRT-PCR was performed in 384-well plates using the Power SYBR Green Master Mix (ThermoFisher Scientific) in a total reaction volume of 10 μl, using a QuantStudio 12K Flex qPCR machine (ThermoFisher Scientific). Following cycling conditions were applied: 2 min at 50 °C, 10 min at 95 °C, 40 x 15 sec at 95 °C and 1 min at 60 °C. Primers are listed on Supplementary Table 1c.

### Bulk mRNA-seq library construction and sequencing

Transcriptome analysis of undifferentiated iPSCs (day 0), sorted NKX2-1 eGFP+ and NKX2-1 eGFP-(day 15) and DCI(±CS)-treated LOs (day 35) was carried out by using the QuantSeq 3′ mRNA-Seq Library Prep Kit for Illumina (REV) with Custom Sequencing Primer (Lexogen) according to manufacturer’s instructions. The 3′ mRNA sequencing libraries were prepared from 110 ng of total input RNA per sample which was isolated with the RNeasy Mini (QIAGEN). Quality of the libraries was evaluated on a Agilent 2100 Bioanalyzer using the High Sensitivity DNA Kit (Agilent Technologies). Samples were sequenced using HiSeq2500 machine.

Differential expression analysis and principal components analysis was performed using DESeq2. GO terms analysis was performed using Panther (Thomas et al., 2003) and STRING platforms (Szklarczyk et al., 2019).

### Single cell RNA-seq (Droplet-sequencing)

#### Generation of single-cell suspensions

Daily sampling of cells during differentiation was performed by detaching the cells from the tissue culture plate using Accutase. Cells where thereafter centrifuged for 5 min at 300 × *g* (4 °C), counted using a Neubauer chamber and critically assessed for single-cell separation and viability. A total of 250,000 cells were aliquoted in 2.5 ml of PBS supplemented with 0.04% of bovine serum albumin (BSA) and loaded for DropSeq at a final concentration of 100 cells/μl.

#### Single-cell RNA sequencing

Drop-seq experiments were performed according to the original Drop-seq protocol (Macosko et al., 2015; Ziegenhain et al., 2017). Using a microfluidic polydimethylsiloxane device (Nanoshift) single cell (100/µl) suspensions were co-encapsulated in droplets with barcoded beads (120b/µl, purchased from ChemGenes Corporation, Wilmington, MA) at rates of 4000 µl/h. Droplet emulsions were collected for 15 min/each prior to droplet breakage by perfluorooctanol (Sigma-Aldrich). After breakage, beads were harvested and the hybridized mRNA transcripts reverse transcribed (Maxima RT, Thermo Fisher). Unused primers were removed by the addition of exonuclease I (New England Biolabs), following which beads were washed, counted, and aliquoted for pre-amplification (2000 beads/reaction, equals ∼100 cells/reaction) using a total of 10 PCR cycles.

#### PCR details

(Smart PCR primer: AAGCAGTGGTATCAACGCAGAGT (100 µM), 2× KAPA HiFi Hotstart Readymix (KAPA Biosystems), cycle conditions: 3 min 95 °C, 4 cycles of 20 s 98 °C, 45 s 65 °C, 3 min 72 °C, followed by 8 cycles of 20 s 98 °C, 20 s 67 °C, 3 min 72 °C, then 5 min at 72 °C) (Macosko et al., 2015).

PCR products of each sample were pooled and purified twice by 0.6× clean-up beads (CleanNA), following the manufacturer’s instructions. Prior to tagmentation, complementary DNA (cDNA) samples were loaded on a DNA High Sensitivity Chip on the 2100 Bioanalyzer (Agilent) to ensure transcript integrity, purity, and amount. For each sample, 1 ng of pre-amplified cDNA from an estimated 600 cells was tagmented by Nextera XT (Illumina) with a custom P5 primer (Integrated DNA Technologies). Single-cell libraries were sequenced in a 100 bp paired-end run on the Illumina HiSeq4000 using 0.2 nM denatured sample and 5% PhiX spike-in. For priming of read 1, 0.5 µM Read1CustSeqB was used (primer sequence: GCCTGTCCGCGGAAGCAGTGGTATCAACGCAGAGTAC).

### Bioinformatics Processing of the data set

The count matrices for each sample were generated by the Drop-seq computational pipeline (version 2.0) as previously described (Macosko et al., 2015). Briefly, STAR (version 2.5.2a) was used for mapping (Dobin et al., 2013). Reads were aligned to the hg19 reference genome (provided by the Drop-seq group, GSE63269). After aggregating the resulting count matrices into one combined object, we used Seurat Version 2.3 (Satija et al., 2015) for downstream analyses. For barcode filtering, we excluded barcodes with less than 400 detected genes. As 1000 cells were expected per sample, the first 1200 cells, sorted by number of transcripts per cell, were used further. A high proportion of transcript counts derived from mitochondria-encoded genes may indicate dying or stressed cells, we removed cells with a percentage of mitochondrial genes of above 20% from downstream analysis. Guided by histograms of quality metrics, further filtering was carried out. Cells with a high number of UMI counts may represent doublets, thus only cells with less than 5000 UMIs were used in downstream analysis.

After the initial filtering of cells, we followed the common preprocessing procedure. The expression matrices were normalized and scaled using Seurat’s NormalizeData() and ScaleData() functions. In order to mitigate the effects of unwanted sources of cell-to-cell variation, cell-cycle effects, percentage of mitochondrial reads and number of counts were regressed out CellCycleScoring() and ScaleData().The 600 top variable genes across the data set were selected via FindVariableGenes() using the default parameters. From this list the known cell cycle genes were excluded. These variable genes were then the basis of the principal component analysis. The first 50 Components were the input for Seurat’s function FindClusters() at a resolution of 2, resulting in 13 clusters. To visualize the clustering result of the high dimensional single-cell data, the UMAP was generated using again 50 components as input for the Seurat function RunUMAP() with a number of neighbors set to 20. To clean up the data set further, a small number of cells disagreeing in the louvain cluster and UMAP embedding were removed. The final meta data and marker gene expression per cluster is reported in Supplementary Table 2 and 3, respectively.

#### Selection of genes with significant association over time

To account for the harsh perturbation in gene expression induced by the medium change, the consequent analyses were split into three stages corresponding to the definite endoderm DE (day 0 to day 6), foregut endoderm FE (day 7 to day 10) and lung progenitors LP (day 11 to day 15). The stage wise temporal analyses were executed using the python package Scanpy (Wolf et al., 2018). The principal components and the neighborhood graph based on the first 10 components was recalculated for each stage separately using Scanpy’s pp.pca() and pp.neighbors(). After generating the diffusion maps via the tl.diffmap() function setting the root cells after manual inspection, the diffusion pseudo time was calculated with tl.dpt(). The diffusion pseudo time for the whole data set was then set by adding the stage-wise pseudo times and scaled to a value between 0 and 1. To assess global connectivity, the louvain clusters were used as input for Partition-based graph abstraction (PAGA) tl.paga(). The weighted edges reflect a statistical measure of connectivity between the groups. Edges with a weight < 0.05 are omitted.

For the identification of genes showing significantly changing expression patterns over time, the following approach was applied by the means of the R packages limma (Ritchie et al., 2015) and splines. As we were particularly interested in the expression pattern towards early lung progenitors, we selected only those cells that were positive for either eGFP or NKX2-1 from days 11 to 15 for the calculation and display. Furthermore, as we did not want to incorporate differences in ribosomal derived genes, such genes were excluded from this analysis. We employed a regression model in which splines are used to model non-linear effects of continuous variables to fit the time-course data. For each gene we fit a natural cubic spline with 4 knots while using the time point of extraction as explanatory variable. UMI counts of each cell were included as a covariate in the model to account for differences in library size.

We rank the genes based on their adjusted p-values. As it is non-trivial to interpret p-values across time, we use the adjusted p-value as a ranking for the genes and visualize the top 200 genes in Fig. 2e. The results are reported in Supplementary Table 4.

#### Stage-wise hierarchical clustering of genes

The clustering analysis was done for the endoderm stage (day 0 to day 6) and the combined endoderm and foregut endoderm stage (day 0 to day 10). For this, genes for both versions with interesting expression patterns were again selected as described previously using the regression model based on spline fits. The scaled expression levels of genes with an adjusted p-value of less than 0.005 were the input for hierarchical clustering using the hdist() from the R package stats. The dendrogram tree was cut into 10 clusters, which were manually re-annotated into 6 final groups for each stage based on their average expression per day. The cluster sizes can be different, thus we chose to display the average expression of the 100 genes with the lowest adjusted p-value per cluster in Fig. 3d,h. For the second clustering analyses, we display the average expression and kinetic profiles for day 6 to day 10. The resulting clusters are reported in Supplementary Table 5.

#### Potential branching into lung and non-lung progenitor cells

To identify genes associated with the potential differentiation branches, we used the signatures from the differential gene expression analysis between eGFP labelled cells versus eGFP negative cells in the initial bulk experiment. As variable genes those genes were selected, which showed a log fold change of above 1 and below -1. To reduce the effect of the media change after day 10, we removed genes that were differentially expressed between the FE and LP stage. In order to guide the dimensionality reduction these 1294 genes were used as input for the PCA. UMAPs and diffusion maps were calculated using Scanpy’s workflow with 50 principal components and the number of neighbors set to 10. In the next step we filtered for the two branches of interest, which started in the FE stage and had their endpoint in the LP stage. The cells were scored to their similarity to the bulk signature via tl.score(), to evaluate if the two potential cell fate trajectories correspond to the branches that we found.

To identify genes showing temporally altered expression patterns along these two potential branches, which are also significantly different across the branches, we employed the python package diffxpy (https://github.com/theislab/diffxpy). The starting population for the two sub-branches (lung and hepatocyte progenitors) was chosen as all cells from FE stage for both trajectories. The cells from the DE stage were divided based on their louvain clusters after visual inspection of the diffusion map and known lineage marker gene expression. The diffusion pseudotime was calculated on these two trajectories separately. Diffxpy’s test.continuous_1d() function runs the test with a spline basis to allow smooth trends. This function was employed on the combination of these two trajectories, using pseudotime as a continuous covariate and a categorical annotation “trajectory” (implying which branch each cell was assigned to) as factor to test for. For visualization purposes, the trajectory-wise pseudotimes were manually binned into “source” and 4 groups each. Average expression of cells per bin is shown for top 100 genes ranked by adjusted p-value from the diffxpy result table in Fig. 4e. The results are reported in Supplementary Table 6.

#### Pairwise gene expression profiles along pseudotime

Cells ordered by diffusion pseudotime were partitioned into 9 bins for the liver and 10 bins for the lung branch similarly as described above. Pairwise gene correlations across the two trajectories were calculated with the cor() function (default parameters, R version 3.6.1) using the average expression in each bin, and visualized using corrplot package in R (version 0.84).

