## Supplementary material for "Single cell trajectory analysis of human pluripotent stem cells differentiating towards lung and hepatocyte progenitors": Table S1 List and catalogue numbers of reagents

**Table 7a. Differentiation experiments**

| **Growth media and supplements** | **Source** | **Cat. No.** |
| --- | --- | --- |
| Activin A | R&D Systems | 338-AC-050 |
| Ascorbic Acid | Sigma Aldrich | A8960-5G |
| BMP4 | R&D Systems | 314-BP-010 |
| B-27 | ThermoFisher Scientific | 17504044 |
| BSA | ThermoFisher Scientific | 15260037 |
| cAMP | Sigma Aldrich | B5386 |
| CHIR99021 | R&D Systems | 4953/50 |
| DAPT | TOCRIS Bioscience | 2634 |
| Dexamethasone | Sigma Aldrich | D4902 |
| DMEM/F12 | ThermoFisher Scientific | 11320033 |
| Dorsomorphin | TOCRIS Bioscience | 3093 |
| FGF-7 (KGF) | R&D Systems | 251-KG-010 |
| FGF-10 | R&D Systems | 345-FG |
| GlutaMAX | ThermoFisher Scientific | 35050038 |
| Ham’s F-12 | ThermoFisher Scientific | 21765029 |
| IBMX | Sigma Aldrich | I5879 |
| IMDM | ThermoFisher Scientific | 12440053 |
| 1-Thioglycerol | Sigma Aldrich | M6145 |
| Growth factor–reduced Matrigel | Corning | 354230 |
| N-2 | ThermoFisher Scientific | 17502048 |
| Penicillin-Streptomycin | ThermoFisher Scientific | 15070063 |
| PneumaCult™-ALI Medium with 6.5 mm Transwell^®^ +Inserts | STEMCELL Technologies | 05022 |
| Retinoic Acid | Sigma Aldrich | R2625 |
| RPMI1640 | ThermoFisher Scientific | 21875034 |
| SB431542 | Miltenyi Biotec | 130-106-543 |
| SHH | R&D Systems | 1845-SH |
| Sodium Butyrate | Sigma Aldrich | B5887-1G |
| Y-27632 | R&D Systems | 1254/10 |

**Table 7b. Antibodies (Immunofluorescence staining and Flow Cytometry)**

| **Antigen** | **Species** | **Manufacturer** | **Cat. No.** |
| --- | --- | --- | --- |

**Primary antibodies**

| FOXA2 | Mouse | Abcam | ab60721 |
| --- | --- | --- | --- |
| SFTPC | Rabbit | Merck Millipore | AB3786 |
| SFTPB | Rabbit | Abcam | ab40876 |
| SCGB1A1 | Mouse | Santa Cruz | sc-365992 |
| MUC5AC | Mouse | Abcam | ab79082 |
| AFP | Mouse | Sigma Aldrich | WH0000174M1-100UG |

**Conjugated antibodies**

| EPCAM-APC | Mouse | BD Biosciences | BDB347200 |
| --- | --- | --- | --- |
| cKIT-APC | Mouse | ThermoFisher Scientific | CD11705 |
| CXCR4-PE | Mouse | ThermoFisher Scientific | MHCXCR404 |

**Secondary antibodies and Isotype controls**

| goat anti-mouse Alexa Fluor 488 | | ThermoFisher Scientific | A-11001 |
| --- | --- | --- | --- |
| goat anti-rabbit Alexa Fluor 647 | | ThermoFisher Scientific | A-21244 |
| goat anti-rabbit Alexa Fluor 546 | | ThermoFisher Scientific | A-11035 |
| donkey anti-mouse Alexa Fluor 594 | | ThermoFisher Scientific | A-21203 |
| mouse IgG2a kappa isotype control, PE | | eBioscience | 12-4724-41 |
| mouse IgG2a kappa isotype control, APC | | eBioscience | 17-4724-81 |

**Table 7c. Primers for SybrGreen qPCR**

| **Gene** | **Sequence** | **Size (bp)** |
| --- | --- | --- |
| *FOXA2* | F: GGGAGCGGTGAAGATGGA  R: TCATGTTGCTCACGGAGGAGTA | 89 |
| *SOX17* | F: GGCGCAGCAGAATCCAGA  R: CCACGACTTGCCCAGCAT | 61 |
| *CXCR4* | F: CACCGCATCTGGAGAACCA  R:GCCCATTTCCTCGGTGTAGTT | 79 |
| *NKX2-1* | F: CTTCCCCGCCATCTCCCGCTTC  R: GCCGACAGGTACTTCTGTTGCTTG | 201 |
| *PAX8* | F: ACTACAAACGCCAGAACCCTACCA  R: CCGGATGATTCTATTAATGGAG | 119 |
| *PAX6* | F: GCGGAGTTATGATACCTACACC  R: GAAATGAGTCCTGTTGAAGTGG | 96 |
| *MUC5AC* | F: GCCTACGAGGATTTTAACAT  R: CAGGACCGGGTGGCCGTTGA | 141 |
| *SFTPC* | F: GTTCTGGAGATGAGCATTGGG  R: GCGATCAGCAGCTGCTGGTAGT | 134 |
| *SFTPB* | F: CTGGCCCAAGGTCGCCGA  R: TGGAGCATTGCCTGTGGTATGG | 121 |
| *SCGB1A1* | F: ATGGACACACCCTCCAGTTATG  R: TGGGCTATTTTTTCCATGAGC | 152 |
| *AFP* | F: GCTTACACAAAGAAAGCCC  R: TAATAATGTCAGCCGCTCC | 142 |
| *GAPDH* | F: GCTCATTTCCTGGTATGACAACG  R: GAGATTCAGTGTGGTGGGGG | 184 |
